# Role of *Cnot6l* in maternal mRNA turnover

**DOI:** 10.1101/362145

**Authors:** Filip Horvat, Helena Fulka, Radek Jankele, Radek Malik, Ma Jun, Katerina Solcova, Radislav Sedlacek, Kristian Vlahovicek, Richard M. Schultz, Petr Svoboda

**Author notes:** Summary blurb: Mice lacking *Cnot6l*, a deadenylase component of the CCR4-NOT complex, are viable but females have ~40% smaller litters. *Cnot6l* is a maternal-effect gene acting in maternal mRNA degradation. These three authors contributed equally to this work (Filip Horvat, Helena Fulka, Radek Jankele). Corresponding author Address for correspondence: Petr Svoboda Institute of Molecular Genetics Videnska 1083 14220 Praha 4 Czech Republic www:http://www.img.cas.cz.

## Abstract

Removal of poly(A) tail is an important mechanism controlling eukaryotic mRNA turnover. The major eukaryotic deadenylase complex CCR4-NOT contains two deadenylase components, CCR4 and CAF1 for which mammalian CCR4 is encoded by *Cnot6* or *Cnot6l* paralogs. We show that *Cnot6l* apparently supplies the majority of CCR4 in the maternal CCR4-NOT complex in mouse, hamster, and bovine oocytes. Deletion of *Cnot6l* yielded viable mice but *Cnot6l*^-/-^ females exhibited ~40% smaller litter size. The main onset of the phenotype was post-zygotic: fertilized *Cnot6l*^-/-^ eggs developed slower and arrested more frequently than *Cnot6l*^+/-^ eggs suggesting that maternal CNOT6L is necessary for accurate oocyte-to-embryo transition (OET). Transcriptome analysis revealed major transcriptome changes in *Cnot6l*^-/-^ ovulated eggs and 1-cell zygotes. In contrast, minimal transcriptome changes in preovulatory *Cnot6l*^-/-^ oocytes were consistent with reported *Cnot6l* mRNA dormancy. A minimal overlap between transcripts sensitive to decapping inhibition and *Cnot6l* loss suggests that decapping and CNOT6L-mediated deadenylation selectively target distinct subsets of mRNAs during OET in mouse.

## INTRODUCTION

During the oocyte-to-embryo transition (OET), maternal mRNAs deposited in the oocyte are gradually replaced by zygotic mRNAs. Consequently, control of mRNA stability is a principal mechanism assuring correct gene expression reprogramming at the beginning of development. Maternal mRNA degradation during mouse OET occurs in several distinct waves (reviewed in detail in [1,2]). Control of mRNA stability involves various mechanisms, many employing protein interaction with the 3’ untranslated region that ultimately targets the terminal 5’ cap and 3’ poly(A) tail structures (reviewed in [3]). The main mammalian mRNA decay pathway involves deadenylation coupled with decapping [4]. Eukaryotic cells employ three main deadenylases: CCR4-NOT complex, PAN2/3 complex, and PARN, which differ in sensitivity to cap structure, poly(A) tail length, and poly(A)-binding protein (reviewed in [5,6]). Cytoplasmic mRNA decay in mammalian cells initiates at the 3’ end and involves sequential deadenylation, first by PAN2/3 followed by CCR4-NOT [4,7]. However, recent data suggest that CCR4-NOT-mediated deadenylation is the main pathway in general mRNA turnover [8].

The multiprotein CCR4-NOT complex (reviewed in [9,10]) was first identified in *Saccharomyces cerevisiae* as a gene regulating glucose-repressible alcohol dehydrogenase 2 (CCR4-NOT stands for <u>C</u>arbon <u>C</u>atabolite <u>R</u>epression <u>4</u>-<u>N</u>egative <u>O</u>n <u>T</u>ATA-less). The mammalian CCR4-NOT complex (Fig. 1A) is composed of a docking platform CNOT1 that binds regulatory components (CNOT2, CNOT3, CNOT4, CNOT9, CNOT10 and CNOT11) and two deadenylase components equivalent to yeast’s CAF1 and CCR4 deadenylases. CAF and CCR4 differ with respect to their relationship with the poly(A)-binding protein (PABP); CCR4 can degrade poly(A) bound with PABP whereas CAF1 degrades free poly(A) [8]. Mammals utilize two paralogs of CAF1 (CNOT7 and CNOT8) and two of CCR4 (CNOT6 and CNOT6L). Thus, a CCR4-NOT complex carries one of four possible combinations of CAF1 and CCR4 homologs. However, the significance of different CCR4-NOT variants remains unclear.

**Figure 1.**
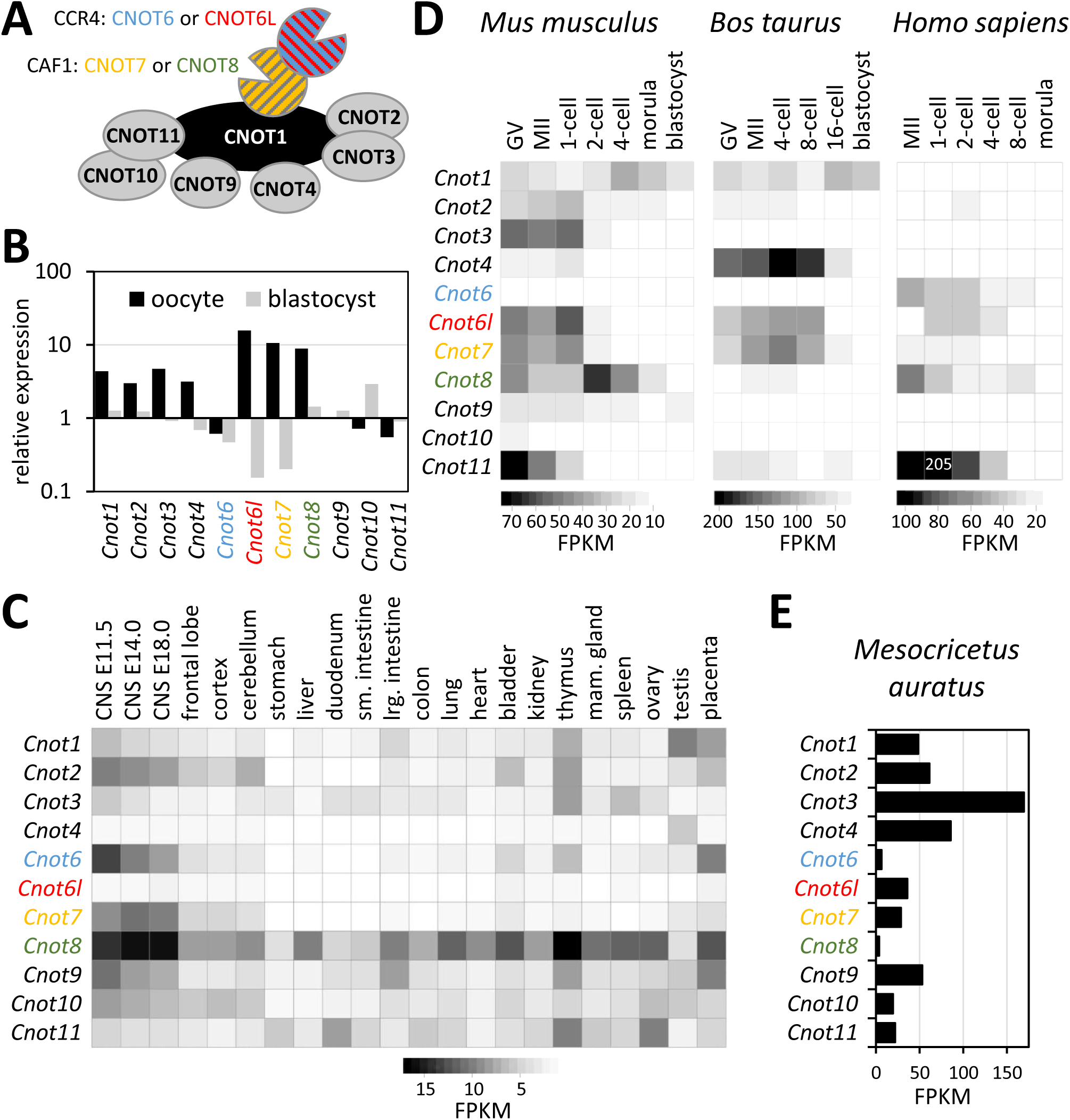
-High maternal expression of *Cnot6l*, an active component of the CCR4-NOT deadenylase complex. **(A)** Schematic depiction of the mammalian CCR4-NOT deadenylase complex. The organization of the complex was compiled from the literature [9,10,41]. Each CCR4-NOT complex contains two active deadenylase proteins, which can make four possible combinations: CNOT7+CNOT6, CNOT7+CNOT6L, CNOT8+CNOT6, and CNOT8+CNOT6L. CNOT6, 6L, 7, and 8 are color-coded for easier navigation in other panels of Figure 1. **(B)** *Cnot6l* transcript is highly enriched in the oocyte relative to somatic cells whereas its alternating paralog *Cnot6* is relatively depleted. The graph shows an expression ratio of CCR4-NOT complex components in oocytes and blastocysts relative to median expression values in somatic tissues calculated from the BioGPS GNF1M.gcrma tissue expression dataset [18] where expression in 61 mouse tissues was set to one. Expression of *Cnot* genes in somatic tissues **(C)**, during oocyte-to-embryo transition in mice, cattle, and humans **(D)**, and in hamster oocytes **(E)**. Heatmaps and the graph show fragments per kilobase per million (FPKMs). Expression data from 22 tissues were selected from the ENCODE polyA RNA-seq mouse tissue panel (GSE49417, [42]), expression analysis of *Cnot* genes in oocytes and early embryos is based on published datasets from indicated species [19-22,35].

The CCR4-NOT complex can be recruited to mRNA by different BTG/Tob proteins, selective RNA binding proteins such as TTP, or upon miRNA binding through TNRC6A-C proteins (reviewed in [11]). Although miRNA-mediated mRNA degradation is insignificant [12], BTG4 plays a major role in maternal mRNA degradation [13]. Another mechanism of selective mRNA targeting by CCR4-NOT is direct recruitment of the complex through YTHDF2, which binds the N6 adenosine (m^6^A) RNA modification [14]. YTHDF2 was linked to selective elimination of maternal mRNAs during oocyte maturation [15].

Maternal mRNAs in mouse oocytes are unusually stable during the growth phase prior to oocyte maturation, which is accompanied with a transition from mRNA stability to instability (reviewed in [2]). This transition also involves recruitment of dormant maternal mRNAs that were accumulated but not (or poorly) translated during the growth phase. Dormant mRNAs encode components of mRNA degradation pathways [13,16,17] and include DCP1A and DCP2, which are critical components of the decapping complex [16]. Inhibiting the maturation-associated increase in DCP1A and DCP2 results in stabilizing a subset of maternal mRNAs that are normally degraded and affects zygotic genome activation [16]. Dormancy was also shown for BTG4 [13] and components of deadenylase complexes: PAN2 for the PAN2/3 complex and CNOT7 and CNOT6L for the CCR4-NOT complex [17].

Here, we report an analysis of *Cnot6l* function during OET in mice. Transcripts encoding the CCR4-NOT complex are relatively more abundant in mouse oocytes than in the blastocyst or in somatic tissues. *Cnot6l* expression apparently supplies the majority of the CCR4 component of the maternal CCR4-NOT complex in mouse, hamster, and bovine, but not human oocytes. Mice lacking *Cnot6l* are viable and fertile. However, zygotes arising from *Cnot6l*^-/-^ eggs develop slower and more likely developmentally arrest than zygotes from heterozygous eggs. Correspondingly, *Cnot6l*^-/-^ females exhibit by ~40% lower fertility. Consistent with the previous report that *Cnot6l* is a dormant maternal mRNA [17], transcriptome analysis revealed minimal transcriptome changes in *Cnot6l*^-/-^ GV oocytes. Nevertheless, there is a subset of maternal mRNAs that are stabilized during oocyte maturation and after fertilization, suggesting that CNOT6L primarily acts in maternal mRNA degradation during oocyte maturation and in zygotes.

## RESULTS & DISCUSSION

### Mammalian CCR4 paralog CNOT6L is highly expressed in oocytes

Several components of the CCR4-NOT complex have a higher relative expression in oocytes in the gcRMA mouse set in the GNF Symatlas database [18] (Fig.1B). Of the four genes encoding active deadenylase components of the CCR4-NOT complex, the CCR4 paralog *Cnot6* showed slightly lower expression when compared to a panel of somatic tissues, whereas transcript abundance of the *Cnot6l* paralog appeared highly enriched in oocytes. These data suggested that CNOT6L could be the main CCR4 deadenylase component during OET (Fig. 1B). In contrast, *Cnot6* appeared to be highly expressed in many somatic tissues, particularly in embryonic neuronal tissues (Fig. 1C).

For further insight into expression of CCR4-NOT complex during OET, we analyzed transcript levels of individual components in RNA-sequencing (RNA-seq) data from OET in mouse [19,20], cow [21] and human [22] (Fig. 1D). In mouse oocytes, *Cnot6l* mRNA was approximately 10-times higher than *Cnot6*, which is expressed during oocyte growth and is apparently not a dormant maternal mRNA [23]. High *Cnot6l* and low *Cnot6* expression during OET was also observed in cow but not in human where the level of *Cnot6* transcript was higher in MII eggs. A subsequent equalization of *Cnot6* and *Cnot6l* expression in human zygotes (Fig. 1D) could be a consequence of cytoplasmic polyadenylation of dormant *Cnot6l* mRNA, which can manifest as an apparent increase in mRNA level in poly(A) RNA-seq data [24]. High *Cnot6l* and low *Cnot6* expression was also found in GV oocytes of golden hamster suggesting that this difference in expression is a conserved feature in rodents (Fig. 1E). Interestingly, CAF1 paralogs *Cnot7* and *Cnot8* showed more variable patterns, including equal expression of both paralogs in mouse, dominating *Cnot7* in bovine and golden hamster, and dominating *Cnot8* paralog expression in human oocytes.

### *Cnot6l* knock-out is viable but exhibits reduced fertility

Given the dominant maternal expression of *Cnot6l* relative to its paralog *Cnot6*, we decided to examine the role of *Cnot6l* in mice using a TALEN-mediated knock-out. We designed two TALEN pairs, which would induce ~31.3 kb deletion affecting exons 5-12 (Fig. 2A). This deletion, which would eliminate the entire CC4b deadenylase domain and a part of the upstream leucine-rich repeat region, was expected to genetically eliminate the CNOT6L protein. We obtained two founder animals carrying two very similar deletion alleles (Fig. 2B and S1). Interestingly, the second allele (Cnot6L-del5-12b) contained a 25-bp insert apparently derived from mitochondrial DNA.

**Figure 2.**
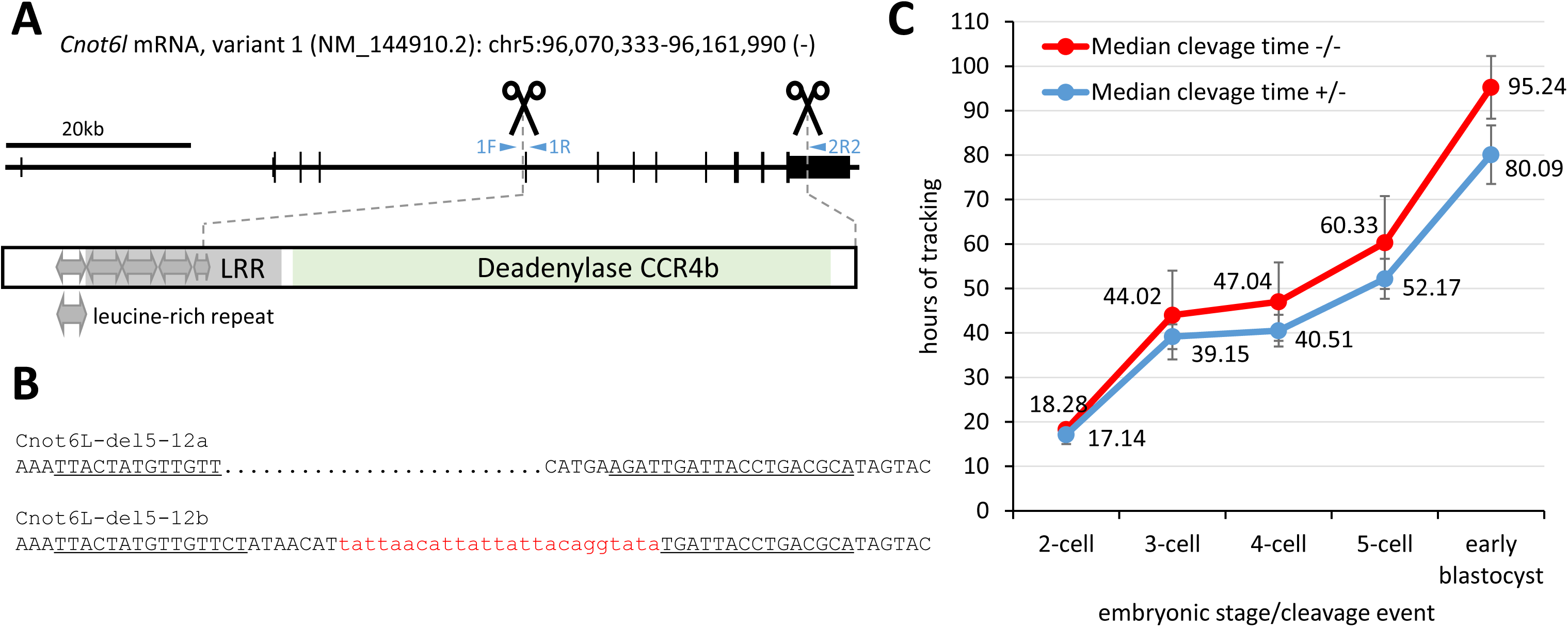
-TALEN-mediated knock-out of *Cnot6l* gene in mice. **(A)** A scheme of *Cnot6l* gene depicting position of the deletion in the genomic DNA and the corresponding part of the CNOT6L protein. Protein domains were mapped using the Conserved Domain Database search [43]. **(B)** Sequences of two alleles identified in F**0** animal, which carry ~31.3 kb deletions of exons 4-11. Underlined sequences indicate TALEN cognate sequences. A short fragment of mitochondrial DNA integrated into the deleted locus is visualized in lower case red font. **(C)** Zygotes from *in vitro* fertilized *Cnot6l*^-/-^ eggs develop significantly slower (t-test, p-value <0.001 for all stages) than zygotes developing from heterozygous eggs. Error bar = S.D. In total, 95 and 133 zygotes produced from *Cnot6l*^-/-^ (-/-) and *Cnot6l*^+/-^ (+/-), respectively, were analyzed using the PrimoVision Time-lapse system. Numbers indicate hours from the point the zygotes were placed into the tracking system (5 h after mixing sperm with COC), which automatically detects first four cleavage events and formation of early blastocysts.

Male and female *Cnot6l*^-/-^ mice appeared normal and were fertile. *Cnot6l* is thus a non-essential gene. Breeding heterozygotes or *Cnot6l*^-/-^ males with *Cnot6l*^+/-^ females yielded on average 6.9 ±1.6 and 6.2 ±1.9 pups per litter, respectively (Table 1), which is consistent with the reported C57BL/6 litter size of 6.2 ±0.2 [25]. We typically observe 6-8 animals per litter in the C57BL/6 strain used to produce mouse models in our facility [20]. Analysis of the *Cnot6l*^-/-^ breeding data showed an average litter size ~4 pups of *Cnot6l*^-/-^ females mated with *Cnot6l*^+/+^, *Cnot6l*^+/-^, or *Cnot6l*^-/-^ males (Table 1). Reduced litter sizes of *Cnot6l*^-/-^ females mated with males of any of the three genotypes were statistically significant (p<0.01, twotailed t-test) when compared to the litter size of *Cnot6l*^+/-^ animals. The breeding data thus indicated a maternal-effect phenotype and showed no evidence for a significant role of zygotic and embryonic expression of *Cnot6l*.

**Table 1.** - Breeding perfomance of *Cnot6l* mutants.

| **F x M** | **+/+** | **+/-** | **-/-** | **n.d.** | **M** | **F** | **litters** | **litter size**<br>**(±SD)** |
| --- | --- | --- | --- | --- | --- | --- | --- | --- |
| +/- x +/- | 26 | 30 | 18 | 2 | 39 | 37 | 11 | 6.9 (±1.6) |
| +/- x -/- | 0 | 22 | 32 | 2 | 28 | 28 | 9 | 6.2 (±1.9) |
| -/- x +/+ | 0 | 48 | 0 | 1 | 27 | 22 | 13 | 3.8 (±1.6) |
| -/- x +/- | 0 | 4 | 11 | 13 | 9 | 19 | 7 | 4.0 (±1.4) |
| -/- x -/- | 0 | 0 | 21 | 0 | 9 | 12 | 5 | 4.2 (±1.8) |

Superovulated knock-out females yielded on average the same number of MII eggs as heterozygote littermates and wild type C57BL/6 (31.5 vs. 32.3 vs. 28.5, respectively, n=6) suggesting that the reduction of litter size occurs during fertilization or post-fertilization. Accordingly, we analyzed early development of zygotes derived from *Cnot6l*^-/-^ and *Cnot6l*^+/-^ eggs fertilized *in vitro* with wild-type sperm and observed a small but significant reduction of fertilization efficiency of *Cnot6l*^-/-^ vs. *Cnot6l*^+/-^ eggs (85 vs. 99%; 133/155 *Cnot6l*^-/-^ vs. 203/205 *Cnot6l*^+/-^ eggs formed zygotes; Fisher’s test p-value <0.001). Analysis of the cleavage times of embryos using a PrimoVision Time-lapse system revealed a small but significant delay in early development that could contribute to the reduced litter size (Fig. 2C). Importantly, although there was no stage-specific arrest of development for *Cnot6l*^-/-^ fertilized eggs, they were two times more likely to fail to reach the blastocyst stage than their *Cnot6l*^+/-^-derived counterparts (38/95 (40.0%) *Cnot6l*^-/-^ vs. 26/133 (19.6%) *Cnot6l*^+/-^ fertilized eggs failed to develop to the blastocyst). This observation suggested reduced developmental competence of *Cnot6l*^-/-^ zygotes and likely accounted for the reduced litter size (Table 1).

### Small but significant transcriptome changes in Cnot6l^-/-^ oocytes and zygotes

To explore the impact of *Cnot6l* loss on the transcriptome during OET, we performed RNA-seq analysis of GV oocytes, MII eggs, and 1-cell zygotes. All replicates showed good reproducibility (Fig. S2 and S3). RNA-seq data showed minimal levels of transcripts arising from the deleted *Cnot6l* allele (Fig. 3A and S4). In addition, we did not observe any compensatory change in *Cnot6* mRNA expression (Fig. 3B).

**Figure 3.**
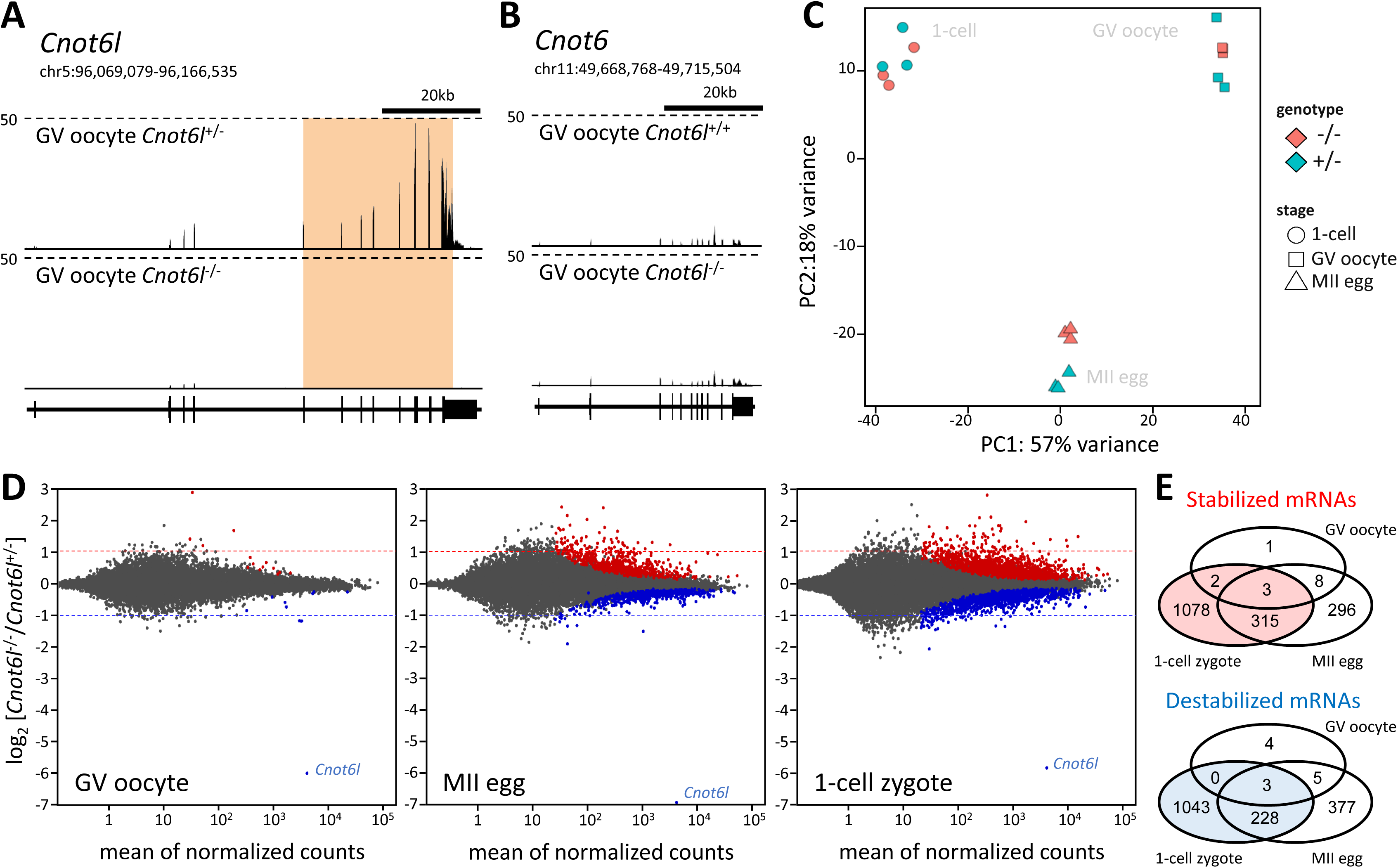
-Transcriptome changes in *Cnot6l* knockout oocytes and zygotes. **(A)** Transcriptional landscape in the *Cnot6l* locus in oocytes from *Cnot6l*^+/-^ and *Cnot6l*^-/-^ animals. Shown is a UCSC Genome Browser snapshot [36] of the *Cnot6l* locus with expression data from one of the replicates of *Cnot6l*^+/-^ and *Cnot6l*^-/-^ samples. The orange region indicates the region deleted in knock-outs. **(B)** Loss of *Cnot6l* expression has no effect on *Cnot6* expression. Shown is a UCSC Genome browser snapshot of the *Cnot6* locus from the same samples as in the panel **(A).(C)** PCA analysis of transcriptomes of *Cnot6l*^-/-^ and control oocytes and zygotes. Heterozygous littermates were used as controls in case of GV and 1-cell zygotes; age-matched C57BL/6 females were used as controls for MII eggs because there were not enough *Cnot6l*^+/-^ littermates for all control samples. **(D)** Differentially expressed trancripts in *Cnot6l*^-/-^ GV oocytes, MII eggs, and 1-cell zygotes. MA-plots depict genes with significantly higher (red) or lower (blue) mRNA abundance. Dashed lines depict 2-fold change for easier navigation. The outlier gene at the bottom of each graph is *Cnot6l*. **(E)** Venn diagrams depicting numbers of genes showing significantly different transcript abundancies in *Cnot6l*^-/-^ oocytes and zygotes.

Transcription from the deleted locus yielded low levels of aberrant transcripts where the splice donor of the third coding exon of *Cnot6l* was spliced with five different downstream splice acceptor sites in the adjacent intron or downstream of the last *Cnot6l* exon (Fig. S4). In all cases, exons spliced with the third coding exon contained stop codons, and thus all transcripts generated from the deleted locus would encode truncated CNOT6L protein composed of the N-terminal leucine repeats. It is unlikely that such isoforms would affect fertility as they would be also present in oocytes of *Cnot6l*^+/-^ females, which have normal fertility.

PCA analysis indicated a small magnitude of changes in knock out samples as samples clustered primarily by developmental stages (Fig. 3C). Analysis of differentially expressed transcripts using DESeq2 package [26] with the default p-value cut-off 0.1 showed minimal transcriptome changes in GV oocytes (only four transcripts showing a significant increase in abundance > 2-fold), which indicates that *Cnot6l* is not required for formation of the maternal transcriptome. There was, however, an apparent progressive transcriptome disturbance in MII eggs and 1-cell zygotes (Fig. 3D). This finding is consistent with the previously reported dormancy of *Cnot6l* [17] and the hypothesis that the reduced litter size of *Cnot6l*^-/-^ females is a maternal-effect phenotype. The magnitude of transcriptome disturbance appears small despite the number of significantly affected genes; if RNA-seq data would be quantified as transcripts per million, higher transcript levels of 622 genes in MII eggs (Fig. 3E) would account for 1.28% of the transcriptome.

Interestingly, the numbers of significantly upregulated and downregulated mRNAs were comparable (Fig. 3E), which was unexpected because transcript stabilization would be the primary expected effect of a deadenylase component loss from the CCR4-NOT complex. It is possible that preventing CNOT6L-mediated deadenylation (hence destabilization) of transcripts from several hundred genes might result in accelerated degradation of other transcripts, noting that we previously observed a similar phenomenon when the maturation-associated increase in DCP1A/DCP2 was inhibited [16]. In any case, when the significantly stabilized transcripts were projected onto transcriptome changes during maturation and following fertilization, it was clear that exclusive CNOT6L-dependent destabilization of maternal transcripts concerns only a smaller fraction of maternal mRNAs degraded during OET (Fig. 4A).

**Figure 4.**
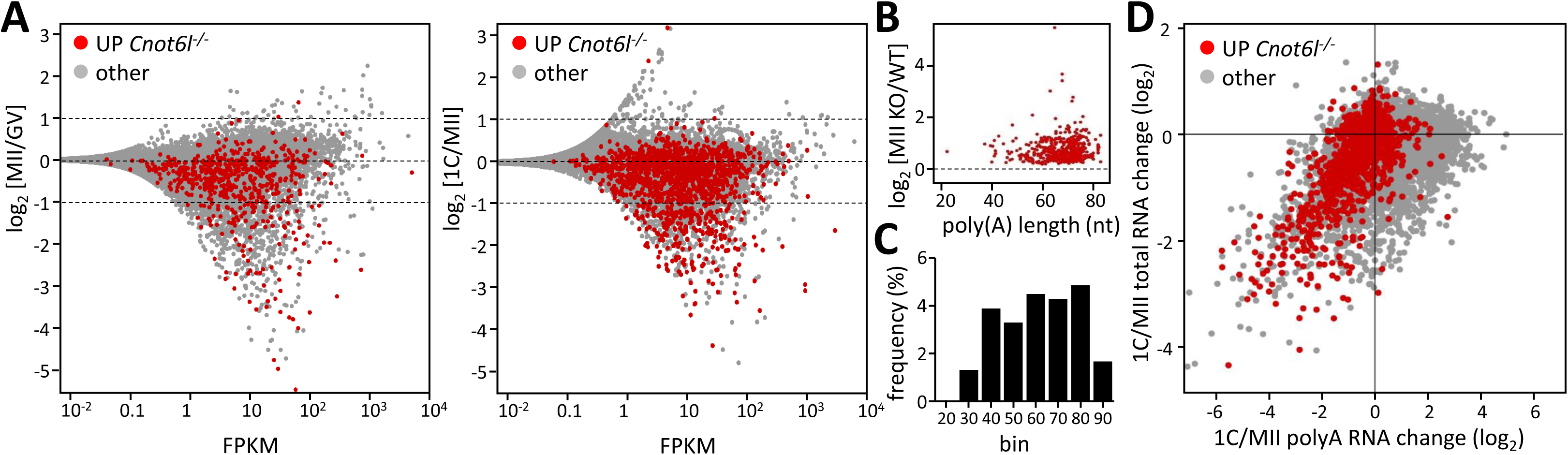
-Transcriptome changes in Cnot6l knockout oocytes and zygotes. **(A)** Projection of differentially expressed trancripts in *Cnot6l*^-/-^ MII eggs and zygotes onto transcripome changes during meiotic maturation and after fertilization. MA-plots were constructed from wild-type samples as indicated and show in red genes with significantly higher mRNA abundance in *Cnot6l*^-/-^ MII eggs (left graph) and zygotes (right graph). **(B)** Analysis of poly(A) tail length of mRNAs significantly increased in *Cnot6l*^-/-^ oocytes during meiosis. The y-axis shows relative upregulation of maternal mRNAs in *Cnot6l*^-/-^ MII eggs (same genes as those labeled in red in MII eggs in Fig, 3D). The x-axis depicts poly(A) tail length in GV oocytes taken from the literature [28]. (C) Relative distribution of mRNAs significantly increased *Cnot6l*^-/-^ oocytes according to poly(A) tail length. RNAs were binned accoring to the poly(A) tail length into ten-nucleotide bins (0-10, 10-20, 20-30, etc.) and the number of transcripts significantly increased in *Cnot6l*^-/-^ MII eggs was divided with the total number of transcripts in each bin. The x-axis numbers represent the upper values of binned poly(A) tail lengths. according to the polyA length in GV oocytes. The y-axis shows relative upregulation of maternal mRNAs in *Cnot6l*^-/-^ MII eggs (correspond to genes labeled in red in MII eggs in Fig, 3D). The x-axis depicts poly(A) tail length in GV oocytes taken from the literature [28]. **(D)** Transcriptome changes in *Cnot6l*^-/-^ zygotes are consistent with a role for CNOT6L in deadenylation during OET. Published RNA-seq data for relative poly(A) RNA and total RNA changes [19,20] were used to construct the plot. In red are shown genes with significantly higher mRNA abundance in *Cnot6l*^-/-^ zygotes. The y-axis shows relative changes of total RNA (i.e., RNA degradation) while the x-axis shows poly(A) RNA changes (i.e., RNA degradation and/or deadenylation).

Relative transcript changes during OET can be problematic to interpret because they may reflect changes in poly(A) tail length and not changes in transcript abundance due to mRNA degradation or transcription [24]. Although the Ovation system used for producing RNA-seq libraries uses total RNA as input material, genome-mapped data show that mRNAs are preferentially sequenced and the sequencing yields a slight bias towards mRNAs with longer poly(A) tails (e.g., *Mos* mRNA, a typical dormant maternal mRNA polyadenylated during meiotic maturation [27], showed an apparent ~17% increased abundance in control wild type MII eggs relative to GV oocytes).

To examine a potential impact of poly(A) tail length on transcript abundance during meiotic maturation, we used a published poly(A) tail sequencing dataset [28] to generate a plot of the relative change in transcript abundance in MII eggs as a function of poly(A) tail length (Fig. 4B). These data showed that transcripts showing relatively increased abundance in *Cnot6l*^-/-^ eggs typically have longer poly(A) tails (60-80 nt). However, when taking into account the distribution of poly(A) lengths in the entire transcriptome, the relative frequency of transcripts with increased abundance in *Cnot6l*^-/-^ eggs was similar for transcripts with poly(A) tails 30 to 80 nt in length (Fig. 4C), suggesting that Fig. 4B data just reflect that most maternal transcripts have poly(A) tails 60-80 nt long.

To further resolve the issue of mRNA abundance vs. poly(A)-length effects in differentially expressed transcripts in zygotes-derived from *Cnot6l*^-/-^ eggs, we used RNA-seq datasets from MII eggs and 1-cell zygotes that were generated from directly-selected poly(A) and from total RNA without any poly(A) bias [19,20]. These data allow distinguishing between true mRNA degradation, which would be observed in the total RNA data, and deadenylation/polyadenylation, which would manifest in the poly(A) data (Fig. 4D). When transcripts showing a significant relative increase in *Cnot6l*^-/-^ zygotes were projected on these data, there was a clear shift to the left on the x-axis, consistent with their deadenylation. Furthermore, a fraction of these transcripts also showed apparent degradation as evidenced by their position on the y-axis (Fig. 4D). Altogether, these data show that maternal *Cnot6l* contributes to maternal mRNA deadenylation and degradation during OET.

Finding that the transcriptome changes following *Cnot6l* loss are restricted to a fraction of deadenylated and degraded maternal mRNAs suggests some selectivity of a CNOT6L-containing CCR4-NOT complex in targeting mRNAs. CCR4-NOT complex recruitment to maternal mRNAs through BTG4 does not appear very selective given the large number of affected maternal mRNAs in *Btg4*^-/-^ MII eggs, which includes ~1/3 of the transcripts showing a relative increase in *Cnot6l*^-/-^ MII eggs (Fig. 5A). These data also suggest that the CAF1 (CNOT7 and CNOT8) component of the CCR4-NOT complex is probably sufficient for a large part of deadenylation mediated by the CCR4-NOT complex. This suggestion is also consistent with the effect of CNOT7 knock-down in early embryos [17], which is apparently more detrimental for early development than the loss of CNOT6L reported here.

**Figure 5.**
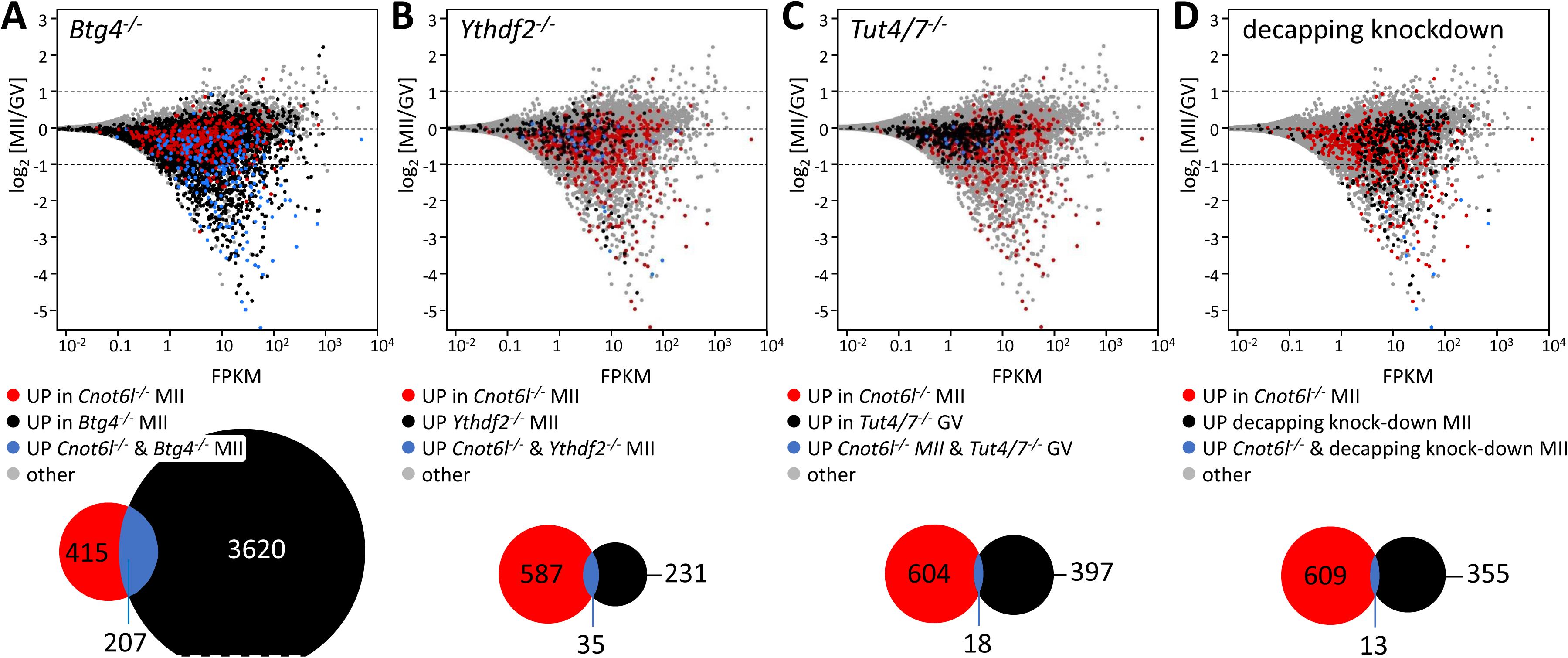
-Comparison of transcriptome changes in *Cnot61*^-/-^ MII eggs with other experimental data. **(A)** Comparison with *Btg4*^-/-^ eggs [13]. **(B)** Comparison with *Ythdf2*^-/-^ eggs [15]. **(C)** Comparison with *Tut4/7*^-/-^ GV oocytes [28]. **(D)** Comparison with eggs lacking production of the decapping complex [16]. In each case, significantly upregulated transcripts in MII eggs were compared with upregulated transcripts in *Cnot6l*^-/-^MII eggs (Fig. 3D). All MA plots were constructed from wild type control replicates from GSE86470.

To gain further insight into potential selectivity of CNOT6L-mediated mRNA deadenylation and degradation, we examined overlaps with transcriptome changes in *Ythdf2* knock-out eggs [15] (Fig. 5B), *Tut4/7*^-/-^ oocytes [28] (Fig. 5C) and in eggs with suppressed decapping [16] (Fig. 5D). In all three cases, transcriptome changes arose during meiotic maturation and concerned hundreds of transcripts. Accordingly, we assessed whether CNOT6L contributes to selective targeting of m^6^A-marked maternal mRNAs during meiotic maturation and to what extent are mRNAs destabilized through CNOT6L and the decapping complex mutually exclusive. In all cases, the overlap of transcripts whose relative abundance is increased in MII eggs was minimal (although statistically significant in the case of the *Ythdf2* knock-out (Fisher’s exact test p-value = 4.06e-14)). Furthermore, transcripts regulated by *Ythdf2* and *Tut4/7* were apparently less-expressed during meiotic maturation (<10 FPKM, Fig. 5B and C) than transcripts targeted by decapping (Fig. 5D).

In any case, maternal mRNAs preferentially targeted through decapping are thus a distinct group from those stabilized upon elimination of *Cnot6l*. This difference becomes apparent when these transcripts are visualized in transcriptome data from unfertilized and fertilized eggs resolved according to relative abundance in total RNA and poly(A) RNA-seq [19,20] (Fig. 6). In this display, the y-axis corresponds to RNA degradation and x-axis reflects poly(A) changes. Deadenylated and degraded RNAs are found in the lower left quadrant. When transcripts upregulated in *Cnot6l*^-/-^ MII eggs or upregulated in MII eggs upon inhibition of decapping are highlighted in this plot, transcripts most sensitive to decapping inhibition seem to be degraded without pronounced deadenylation, unlike transcripts sensitive to *Cnot6l* loss (Fig. 6).

**Figure 6.**
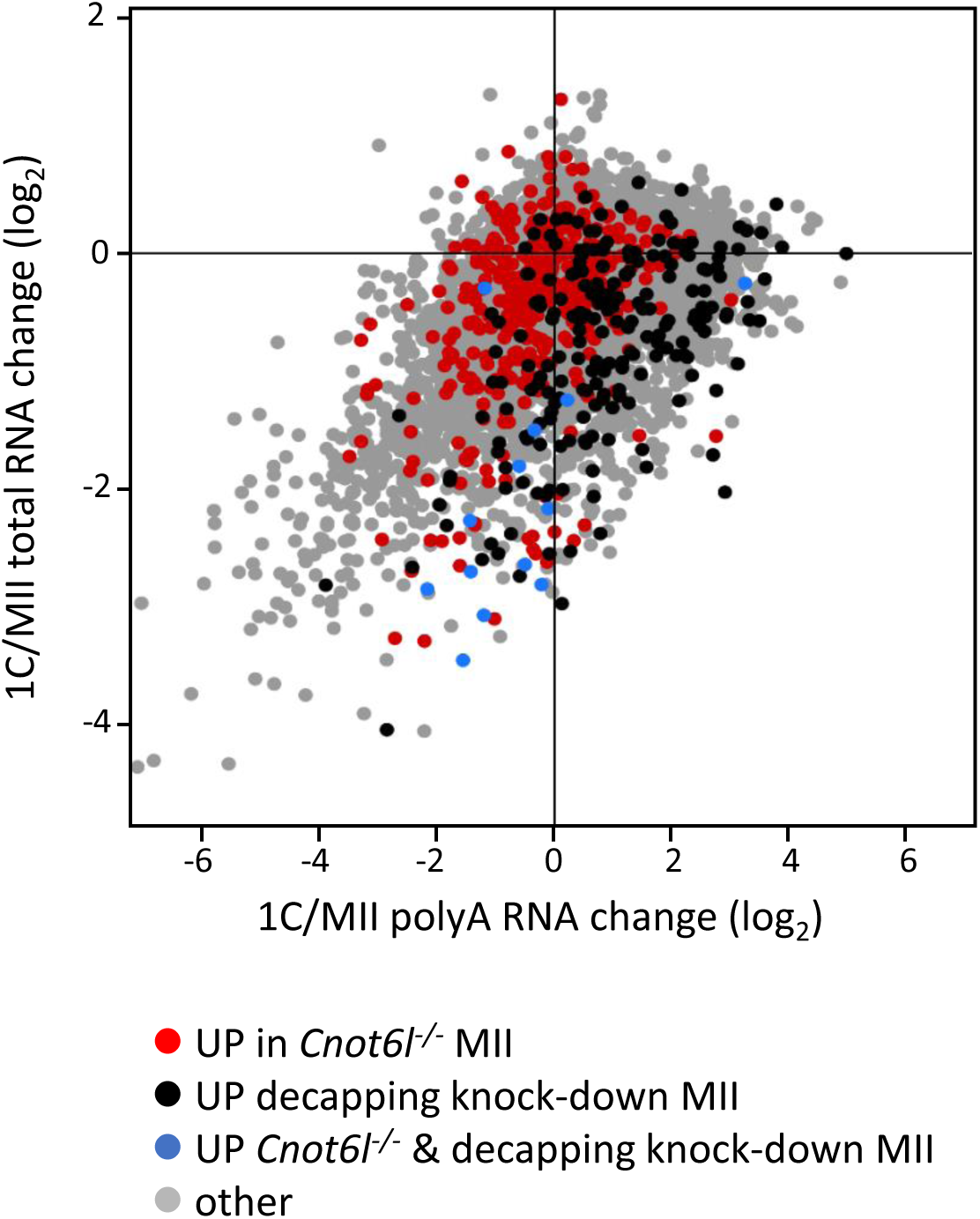
-Comparison of transcriptome changes in *Cnot61*^-/-^ MII eggs with other experimental data. Projection of transcripts with relatively increased abundance in *Cnot6l*^-/-^ MII eggs and eggs with blocked decapping (Fig. 5D) onto transcripome changes in zygotes. The plot was constructed from published data [19,20] as in Fig. 4D. The y-axis shows relative changes of total RNA RNA-seq (i.e., RNA degradation) whereas the x-axis shows poly(A) RNA changes from poly(A) RNA-seq (i.e., RNA degradation and/or deadenylation). In red are shown transcripts with significantly higher mRNA abundance in *Cnot6l*^-/-^ MII eggs. In black are shown transcripts with significantly higher mRNA abundance in MII eggs upon inhibition of decapping. In blue are shown transcripts upregulated in both conditions.

A selective function has been proposed for CNOT6-mediated deadenylation of maternal mRNAs. CNOT6 is present in full-grown GV oocytes in cortical foci and regulates deadenylation of mRNAs such as *Orc6* or *Slbp* that were transiently polyadenylated during early meiotic maturation [23]. Remarkably, CNOT6 and CNOT6L paralogs are highly similar at the protein level (Fig. S5); the major differences concern five amino acid residues longer N-terminus of CNOT6L and five amino acid residues insertion in CNOT6 at the end of the N-terminal leucine-rich repeat region, which stems from using alternative splice donor. Further research should reveal whether these differences underlie any distinct recruitment of CCR4-NOT complexes carrying these paralogs, or whether apparent selectivity is determined by other factors, such as the length of the poly(A) tail, differential expression of the paralogs or their specific localization in oocytes and zygotes.

Taken together, we show that loss of *Cnot6l* in mice results in reduced fertility. Although we cannot rule out that some of the effects observed in the oocyte or early embryos could be indirect effects of a role for *Cnot6l* (e.g., in granulosa cells), the phenotype is presumably a consequence of perturbed deadenylation and degradation of maternal mRNAs during OET. Because *Btg4*^-/-^ eggs exhibit much larger transcriptome changes than *Cnot6l*^-/-^ eggs, CNOT6L-mediated deadenylation appears rather selective. It is presently unclear if this selectivity stems from truly selective targeting (e.g., dependent on recruitment of CNOT6L-containing CCR4-NOT complex directly through CNOT6L). Given a possible redundancy with the CAF1 component of the CNOT6L complex and/or other RNA degrading mechanisms, we speculate that a spectrum of transcripts targeted by the CNOT6L-containing CCR4-NOT complex is much broader and transcripts showing a relative increase upon loss of *Cnot6l* are less targeted by redundant mechanisms.

## MATERIAL AND METHODS

### Oocyte and embryo collection

Oocytes and early embryos were obtained from superovulated mice as described previously [29]. Resumption of meiosis during culture of GV oocytes was prevented with 0.2 mM 3-isobutyl-1-methyl-xanthine (IBMX, Sigma). Animal experiments were approved by the Institutional Animal Use and Care Committees (approval no. 024/2012) and were carried out in accordance with the European Union regulations.

### Production of *Cnot6l* knock-out model

*Cnot6l* knockout mice were produced by the Transgenic and Archiving Module of the Czech Centre for Phenogenomics (http://www.phenogenomics.cz/), Institute of Molecular Genetics ASCR, using using TAL Effector Nucleases (TALENs, reviewed in [30]) designed to delete coding exons 4 −11 (Fig. 2A). TALEN plasmids were produced as described previously [31]. Target sites for TALEN pairs inducing a ~31.3 kb deletion (chr5:96,075,067-96,106,332 (GRCm38/mm10)) were identified using SAPTA tool [32], TALEN off-targeting was addressed with the PROGNOS tool [33]. The following RVD repeats were used to generate individual TALENs:

C6L-1R repeats NH NI HD NI NI NG NG NH HD NI NI NG NG NG HD - (cognate sequence: T0 GACAATTGCAATTTC)

C6L – 1F repeats NG NI HD NG NI NG NH NG NG NH NG NG HD NG - (cognate sequence: T0 TACTATGTTGTTCT)

C6L - 4F repeats NG NI NG NI NG NI HD HD NG HD NI NI NH HD NI NG HD HD - (cognate sequence: T0 TATATACCTCAAGCATCC)

C6L 4R repeats NH HD NH NG HD NI NH NH NG NI NI NG HD NI NI NG HD NG - (cognate sequence: T0 GCGTCAGGTAATCAATCT)

TALEN RNAs for injection were produced as described previously [31]. A sample for microinjection was prepared by mixing all four TALEN RNAs in ultra-pure water at concentration of 4 ng/μl each. This mixture was loaded into injection capillary and injected into male pronuclei of C57BL/6 1-cell embryos.

Genotyping was performed by PCR on lysates from tail biopsies from four weeks-old animals using genotyping primers Cnot6l-1F (5’-GTCATCAGGTTTGGCAGCAAGC-3’) and Cnot6l −1R (5’-CTAAGAAGTGTGTGGTGCATCAGC-3’) for the wild-type allele (yielding a 597 bp product) and Cnot6l-1F and Cnot6l-2R2 (5’-CAGAGAAGAAAGCCCACCCG-3’) for the deletion (yielding a predicted 357 bp product).

### Analysis of preimplantation development

Mice were superovulated and cumulus oocyte complexes (COCs) were isolated as described previously [29]. Sperm of C57BL/6J (8-12 weeks old) males were used for in vitro fertilization (IVF). Males were sacrificed by cervical dislocation and sperm were isolated from cauda epididymis, capacitated for 1h in Human Tubal Fluid (HTF) medium and mixed with COCs in HTF. IVF was performed for 5 h. Next, zygotes containing two pronuclei were selected and preimplantation development was analyzed by the PrimoVision Time-lapse system (Vitrolife) with 15-min acquisition settings (t=0; start of recording). Embryos were cultured in KSOM medium at 37°C under 5% CO_2_ until the blastocysts stage. Automatically recorded times for pre-set cleavage events were confirmed by personal inspection and the median time was plotted against the embryonic stage. The experiment was repeated five times. Heterozygous and homozygous females used in each experiment were littermates, the eggs were fertilized by a single male and embryos developed side-by-side in a single incubator. Representative recorded videos are provided in the supplemental material.

### RNA sequencing

Total RNA was extracted from triplicates of 25 wild type or knock-out GV oocytes, MII eggs or 1-cell zygotes using a PicoPure RNA Isolation Kit with on-column genomic DNA digestion according to the manufacturer’s instructions (Thermo Fisher Scientific, Philadelphia, PA). Each sample was spiked in with 0.2 pg of synthesized *Renilla* luciferase mRNA before extraction as a normalization control. Non-stranded RNAseq libraries were constructed using the Ovation RNA-seq system V2 (NuGEN, San Carlos, CA) followed by Ovation Ultralow Library system (DR Multiplex System, NuGEN, San Carlos, CA). RNAseq libraries were pooled and sequenced by 125 bp paired-end reading using the Illumina HiSeq at the High Throughput Genomics Core Facility at University of Utah, Salt Lake City, UT. RNA-seq data were deposited in the Gene Expression Omnibus database under accession ID GSE86470.

### Bioinformatics analyses

#### Mapping of Illumina RNA-seq reads on the mouse genome

All RNA-seq data were mapped using the STAR mapper [34] version 2.5.3a as described previously [35], except of allowing all multimapping reads:

STAR ‐‐readFilesIn $FILE1 $FILE2 ‐‐genomeDir $GENOME_INDEX ‐‐runThreadN 8 ‐‐genomeLoad LoadAndRemove ‐‐limitBAMsortRAM 20000000000 ‐‐readFilesCommand unpigz -c ‐‐outFileNamePrefix $FILENAME ‐‐outSAMtype BAM SortedByCoordinate ‐‐outReadsUnmapped Fastx ‐‐outFilterMultimapNmax 99999 ‐‐outFilterMismatchNoverLmax 0.2 ‐‐sjdbScore 2

The following genome versions were used for mapping the data: mouse - mm10/ GRCm38, human – hg38/GRCh38, cow – bosTau8/ UMD3.1, hamster - MesAur1.0 (GCF_000349665.1). Annotated gene models for all organisms corresponding to their respective genome versions were downloaded from Ensembl database as GTF files. Only protein coding genes were used in all subsequent analyses. Data were visualized in the UCSC Genome Browser by constructing bigWig tracks using the UCSC tools [36].

#### Differential expression analysis of RNA-seq data

Analysis of genes differentially expressed in knockouts compared to wild type in different developmental stages was done in R software environment. Mapped reads were counted over exons grouped by gene:

GenomicAlignments::summarizeOverlaps(features = exons, reads = bamfiles, mode = “Union”, singleEnd = FALSE, ignore.strand = TRUE)

Statistical significance and fold changes in gene expression were computed using DESeq2 package [26] from RNA-seq data prepared as biological triplicates. Briefly, DESeq2 analysis starts with a matrix of read counts obtained with *summarizeOverlaps()* command above in which each row represents one gene and each column one sample. Read counts are first scaled by a normalization factor to account for differences in sequencing depth between samples. Next, dispersion (i.e., the variability between replicates) is calculated for each gene. Finally, negative binomial generalized linear model (GLM) is fitted for each gene using those estimates and normalized counts. GLM fit returns coefficients indicating the overall expression strength of the gene and coefficients (i.e., log_2_-fold change) between treatment and control (in our analysis knock-out and wild type samples). Significance of coefficients in GLMs are tested with the Wald test. Obtained p-values are adjusted for multiple testing using the Benjamini and Hochberg False Discovery Rate procedure [37]. In our analysis, expression changes with p-adjusted values smaller than 0.1 (the default DESeq2 cutoff) were considered significant.

#### Differential expression analysis of microarray data (decapping complex, YTHDF2)

Microarray data were normalized and background corrected using RMA [38] (*Ythdf2* data) or GC-RMA [39] (decapping complex data) algorithms. Statistical significance and fold changes in gene expression were computed using SAM method [40].

#### PCA plot

Principal component analysis was computed on count data transformed using regularized logarithm (rlog) function from DESeq2 [26] R package.

#### Statistical analysis

Fisher’s exact test was used to evaluate the significance of number of genes showing increased transcript abundance in *Cnot6l*^-/-^ MII eggs and MII eggs with reduced decapping complex or knock-outs of *Ythdf2* or *Btg4.* A level of P < 0.05 was considered to be significant.

## ACKNOWLEDGEMENTS

We thank Vedran Franke and Josef Pasulka for help with data analysis. This research was supported by the Czech Science Foundation (CSF) grant P305/12/G034 and by the Ministry of Education, Youth, and Sports (MEYS) project NPU1 LO1419. Additional support of co-authors included a CSF grant 17-08605S to HF, LM2015040 (Czech Centre for Phenogenomics), CZ.1.05/1.1.00/02.0109 (BIOCEV - Biotechnology and Biomedicine Centre of the Academy of Sciences and Charles University), and CZ.1.05/2.1.00/19.0395 (Higher quality and capacity for transgenic models) support by MEYS and RVO 68378050 by Academy of Sciences of the Czech Republic to RS, the European Structural and Investment Funds grant for the Croatian National Centre of Research Excellence in Personalized Healthcare (contract #KK.01.1.1.01.0010), Croatian National Centre of Research Excellence for Data Science and Advanced Cooperative Systems (contract KK.01.1.1.01.0009) and Croatian Science Foundation (grant IP-2014-09-6400) to KV, and a grant from NIH (HD022681) to RMS.

## AUTHOR CONTRIBUTION

FH – Investigation, Data curation, Software, Formal analysis, Writing – review & editing

HF – Investigation, Formal analysis, Writing – review & editing

RJ – Investigation, Formal analysis, Writing – review & editing

RM – Investigation, Writing – review & editing

MJ – Investigation

KS – Methodology

RS – Resources

KV – Resources, Software, Data curation

RMS – Conceptualization, Funding Acquisition, Project administration, Writing – review & editing

PS – Conceptualization, Funding Acquisition, Project administration, Supervision, Visualization, Writing – original draft

## SUPPLEMENTAL MATERIAL

**Supplemental Figure S1.**
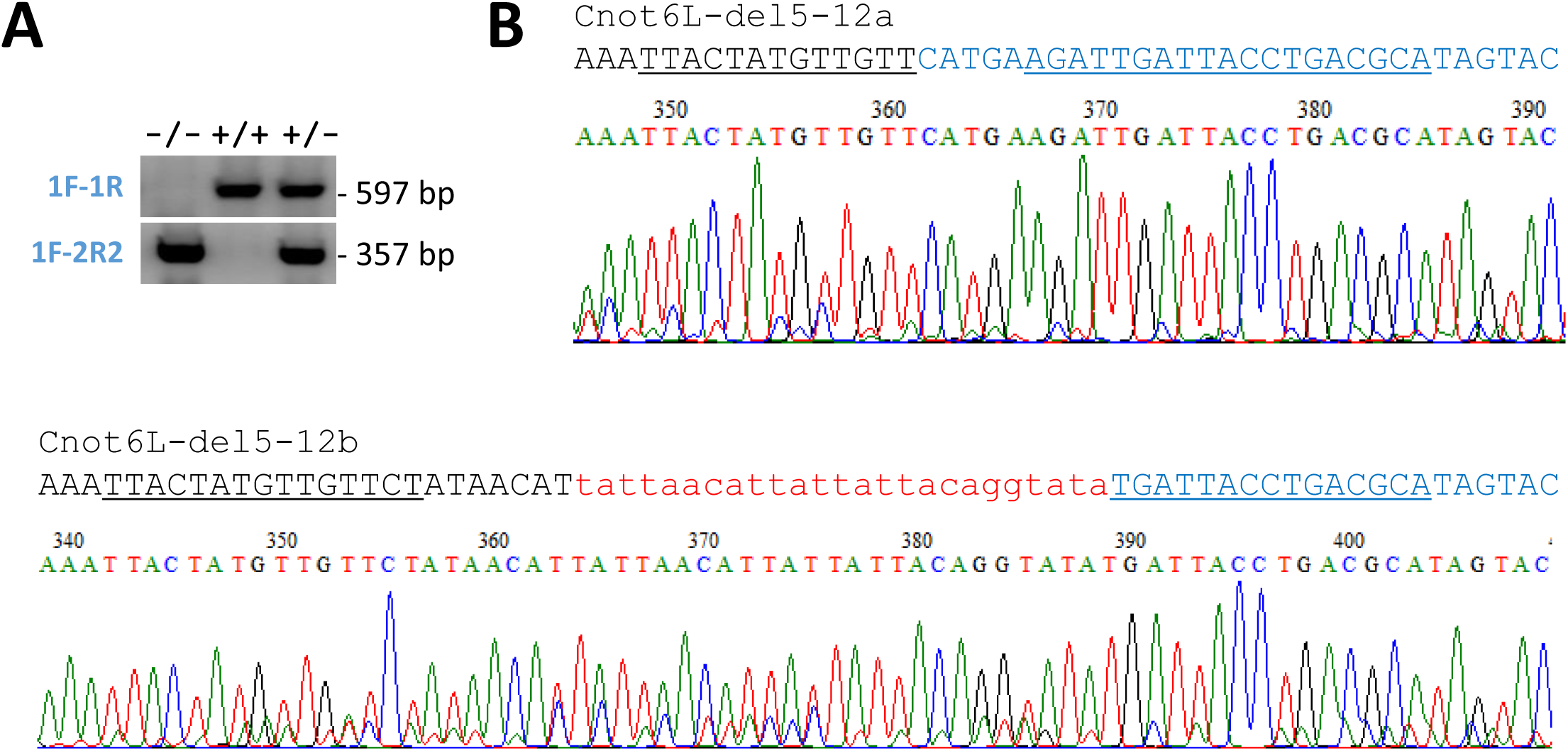
*Cnot6l*^-/-^ mouse genotyping. (A) An example of genotyping F1 mutant mice with three different genotypes. (B) Identification of Cnot6L-del5-12a and Cnot6L-del5-12b alleles in F0 animal by Sanger sequencing of genotyping PCR products. Black and blue letters indicate deletion flanks, the red lower-case letters depict a 25-bp insert matching mitochondrial DNA sequence.

**Supplemental Figure S2.**
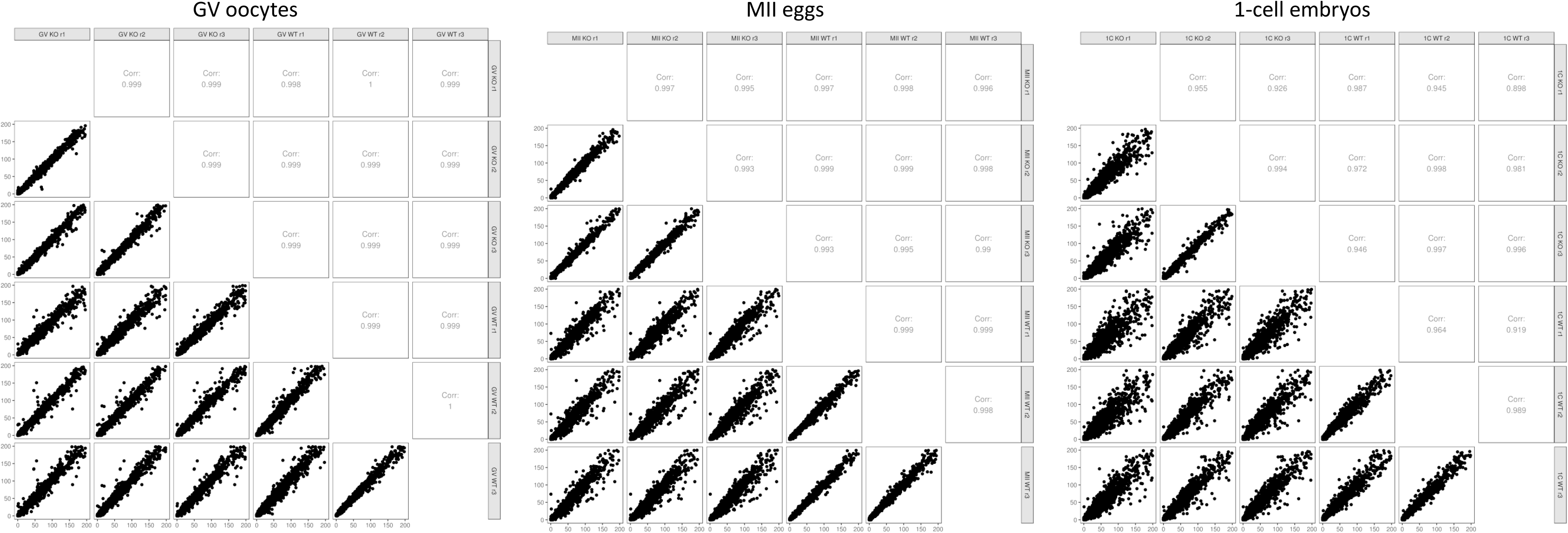
Matrix of scatter plots showing correlation between all samples from each developmental stage. Corresponding correlation coefficients are shown in the upper portion of the matrix.

**Supplemental Figure S3.**
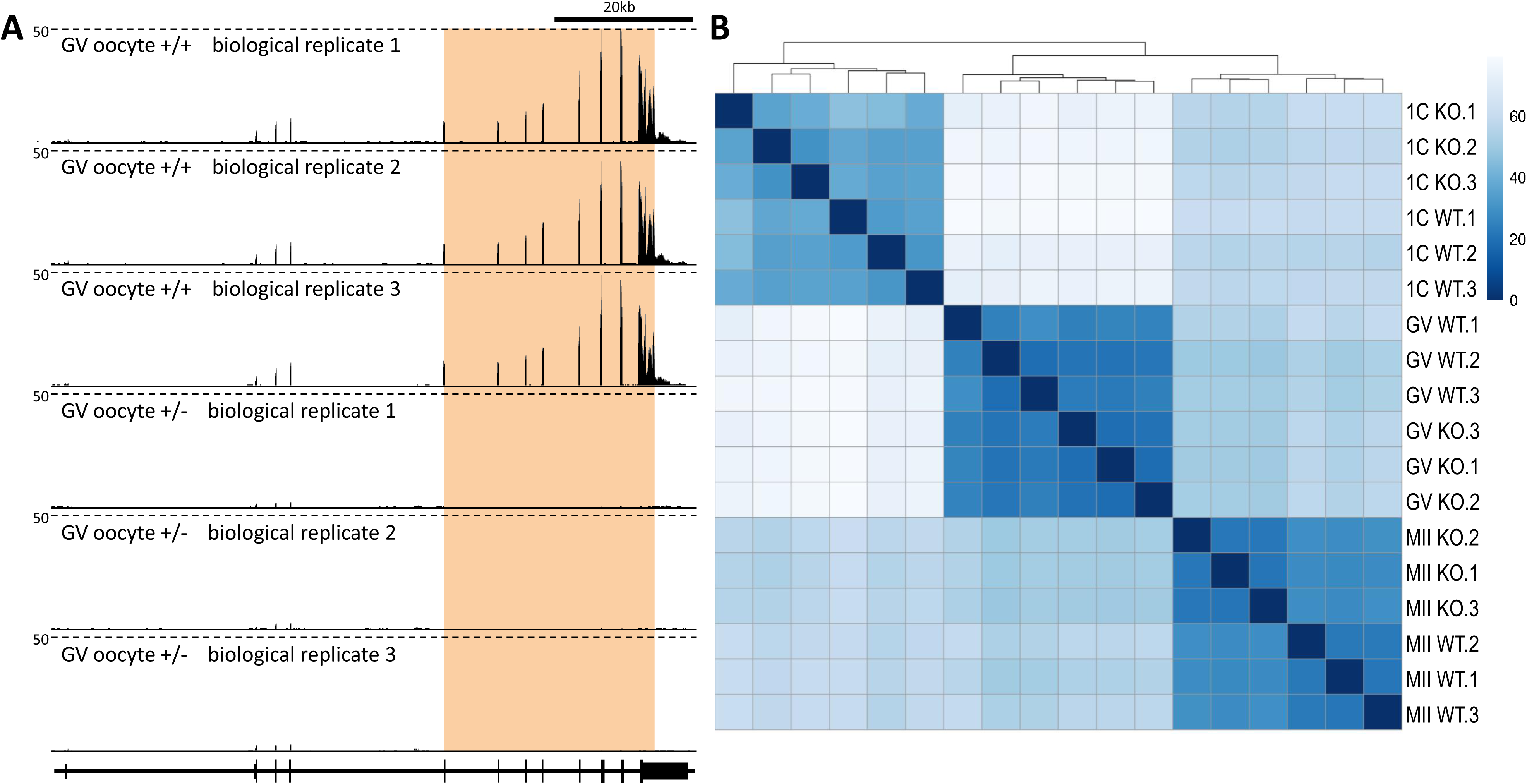
**(A)** Consistent expression profiles in the *Cnot6l* locus in replicate samples. Shown is a UCSC Genome Browser snapshot [36] of transcriptional landscape in the *Cnot6l* locus in GV oocytes from *Cnot6l*^+/-^ and *Cnot6l*^-/-^ animals. The orange region indicates the position of the deletion in knock-outs. The scale (50) depics counts per million (CPM). **(B)** Heatmap showing distances between samples. Euclidean distances between samples were computed on count data transformed using regularized logarithm (rlog) function from DESeq2 [26] R package. Distances were clustered using hierarchical clustering complete linkage method.

**Supplemental Figure S4.**
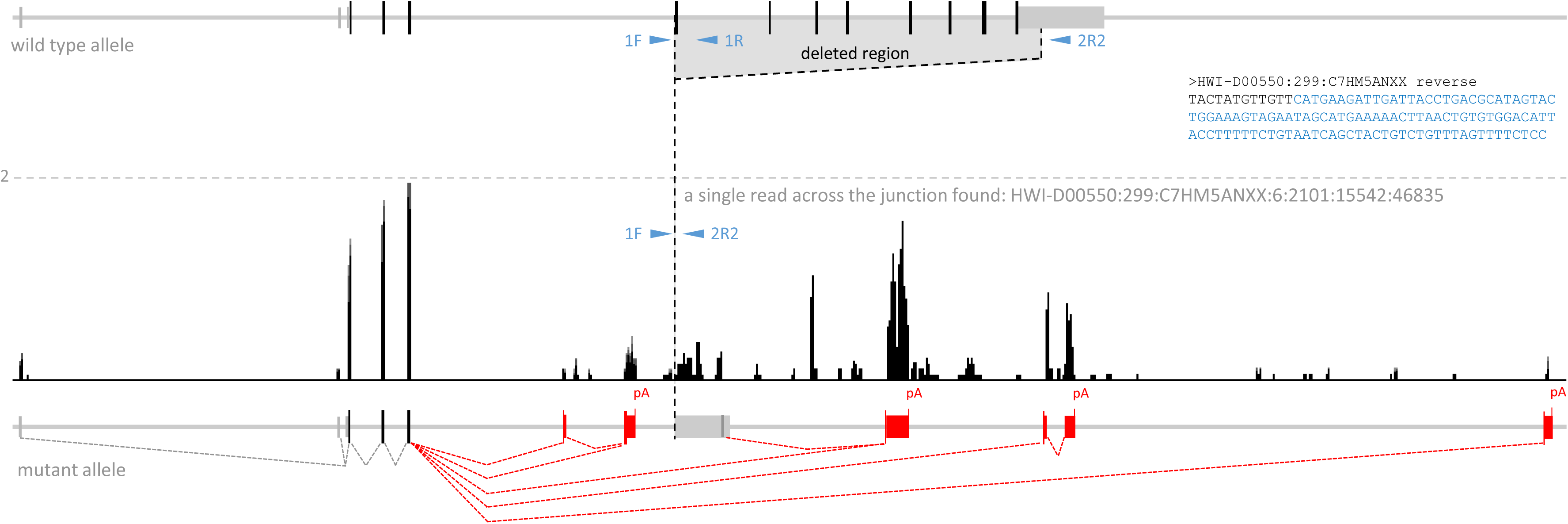
Transcritpional remodelling of the deleted *Cnot6l* locus. Shown is a UCSC Genome Browser snapshot [36] of transcriptional landscape of the mutant *Cnot6l* locus (Cnot6L-del5-12a allele). The dashed depicts the scale of two counts per million (CPM). Also shown is the sequence of a single RNA read found to map across the junction (HWI-D00550:299:C7HM5ANXX). Novel splicing events (indicated with red dashed lines) were identified by analyzing split reads mapping to the locus. The remaining reads mapping to the locus either represent multimappers mapping to repetitive sequences or potential fragments from nascent transcripts. In total, five different splice acceptors for the splice donor in the third coding exon were found. Extension of the *Cnot6l* coding sequence in alternative 3’ end exons is indicated by the height of the exon depiction.

**Supplemental Figure S5.**
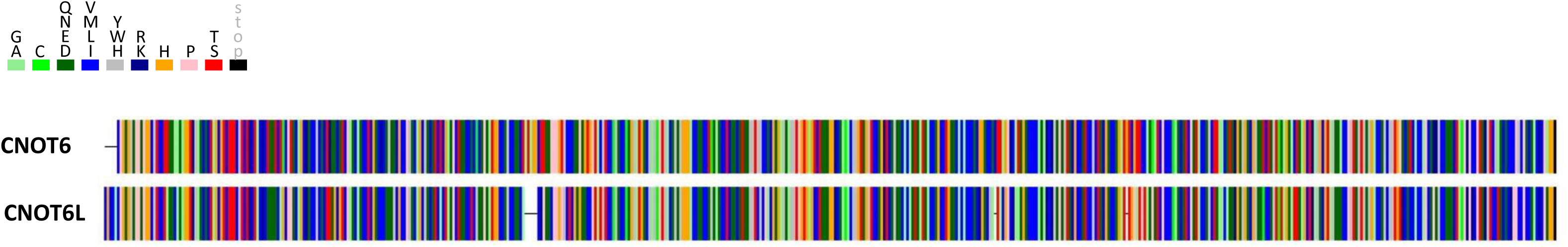
Sequence comparison of CNOT6 and CNOT6L proteins.

**Supplemental Table S1** Differentially expressed genes in *Cnot6l*^-/-^ GV oocytes

**Supplemental Table S2** Differentially expressed genes in *Cnot6l*^-/-^ MII eggs

**Supplemental Table S3** Differentially expressed genes in *Cnot6l*^-/-^ 1-cell embryos

